## Supplementary figures and images for "Genome-Wide Analysis of Horizontal Transfer in Non-Model Wild Species from a Natural Ecosystem Reveals New Insights into Genetic Exchange in Plants"

### Supplemental Figure 1

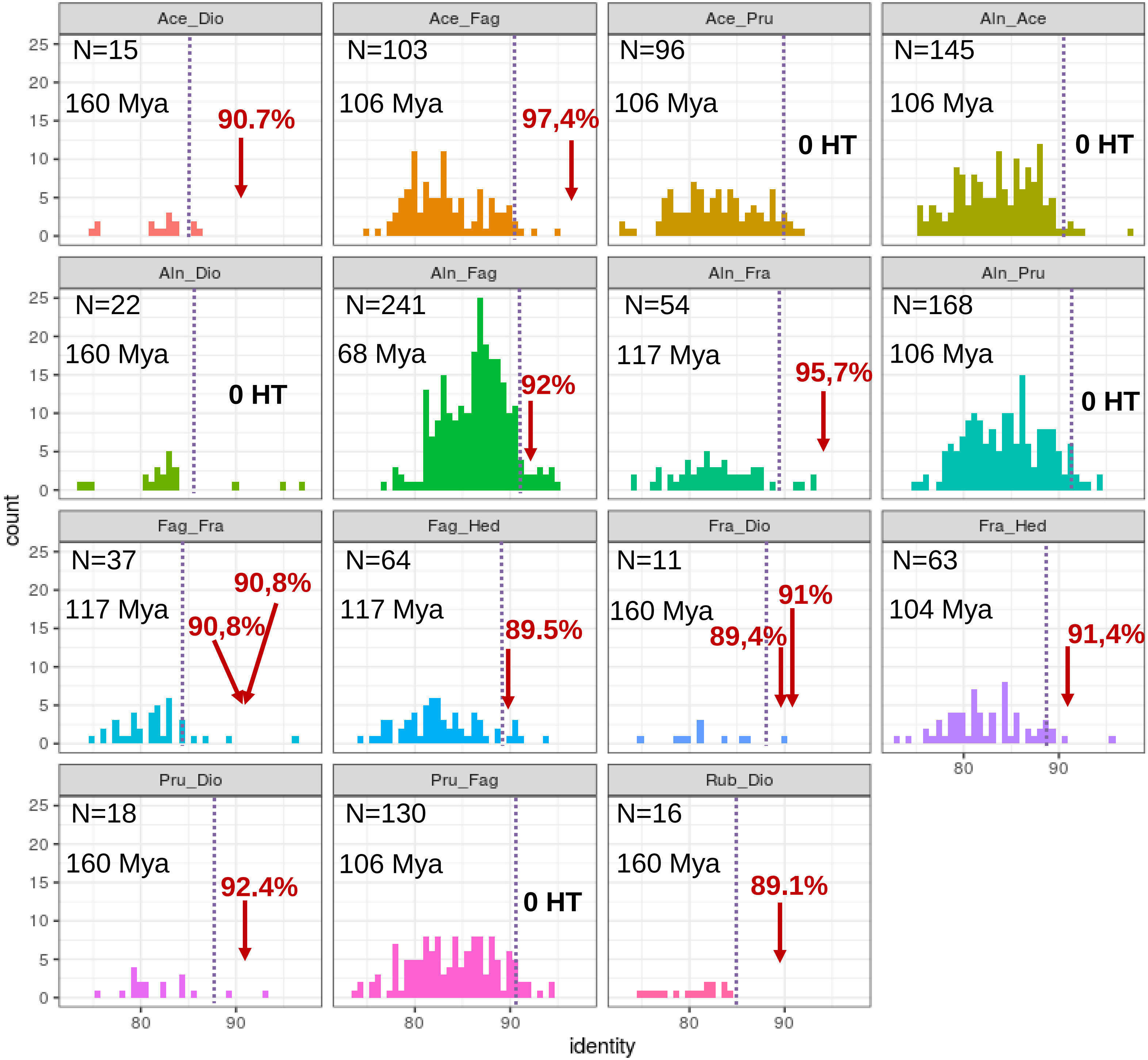

### Supplemental Figure 2

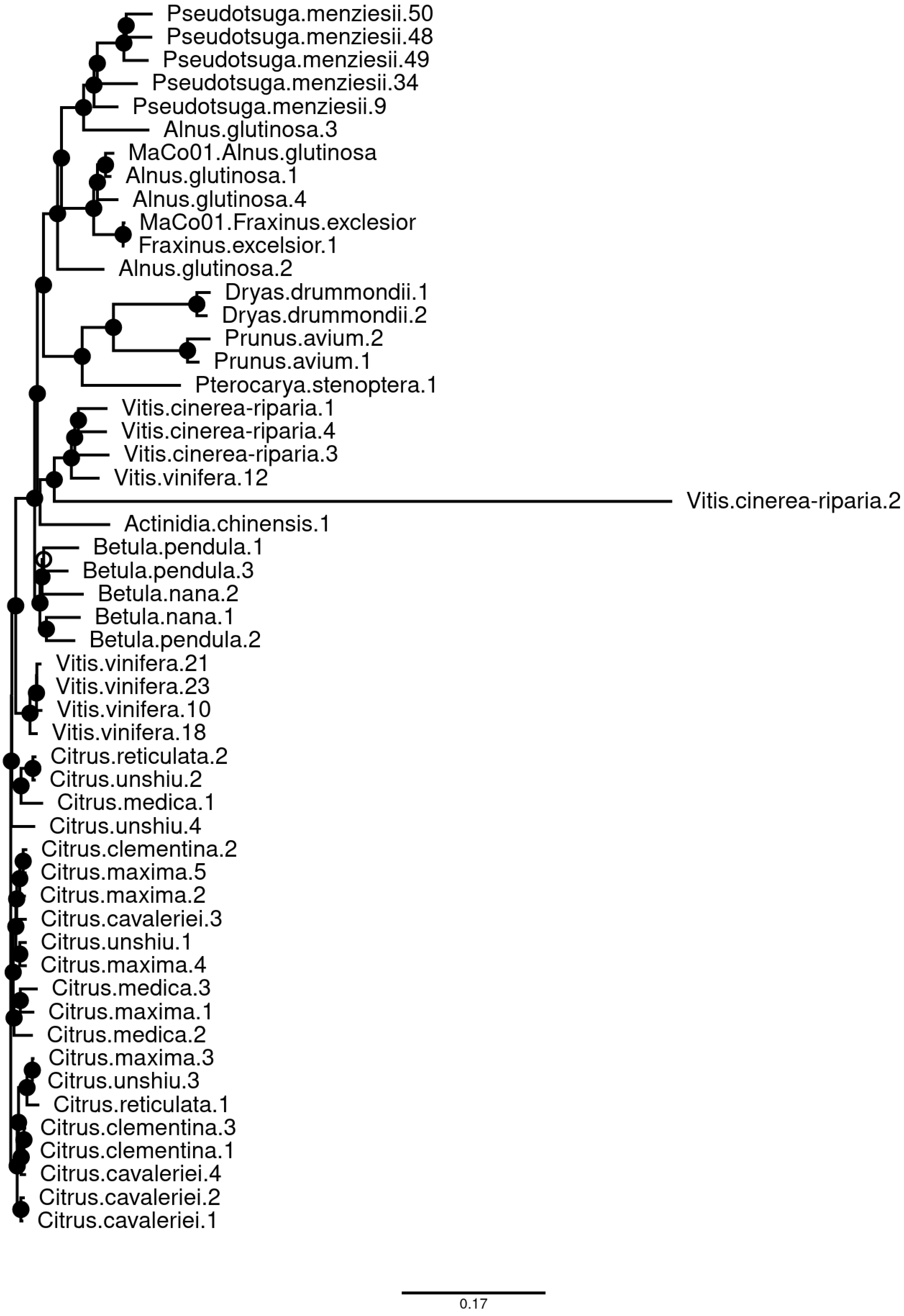

### Supplemental Figure 3

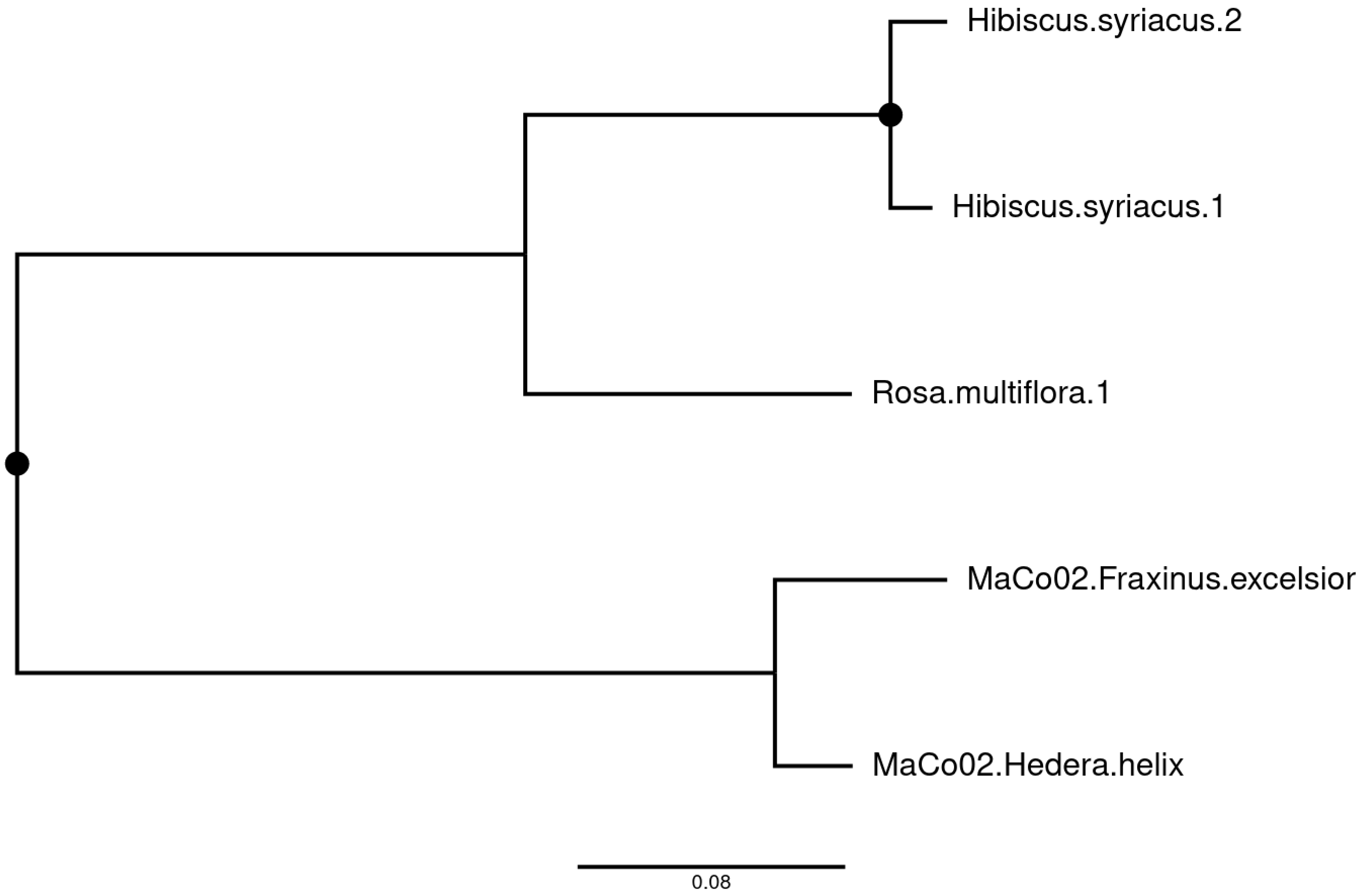

### Supplemental Figure 4

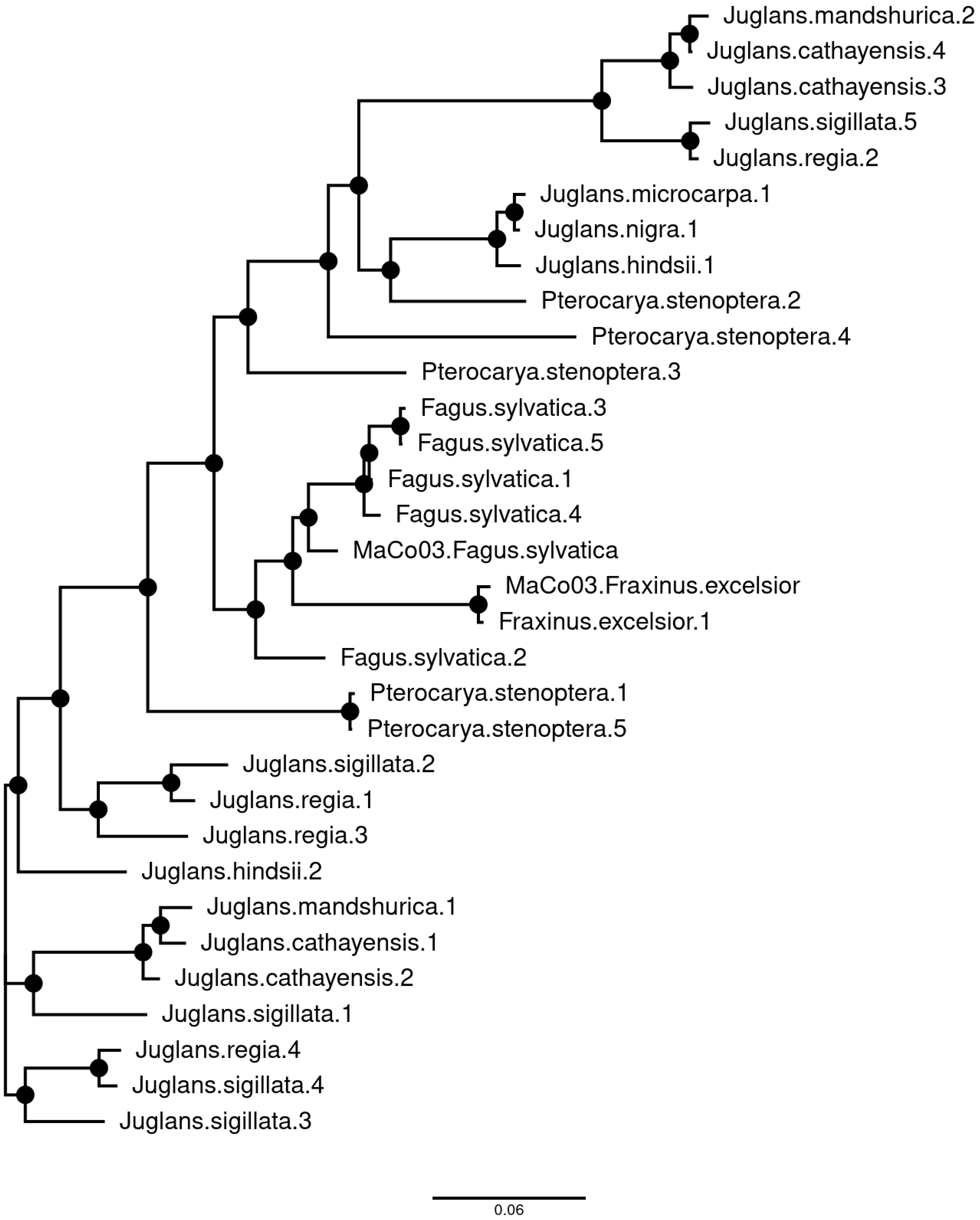

### Supplemental Figure 5

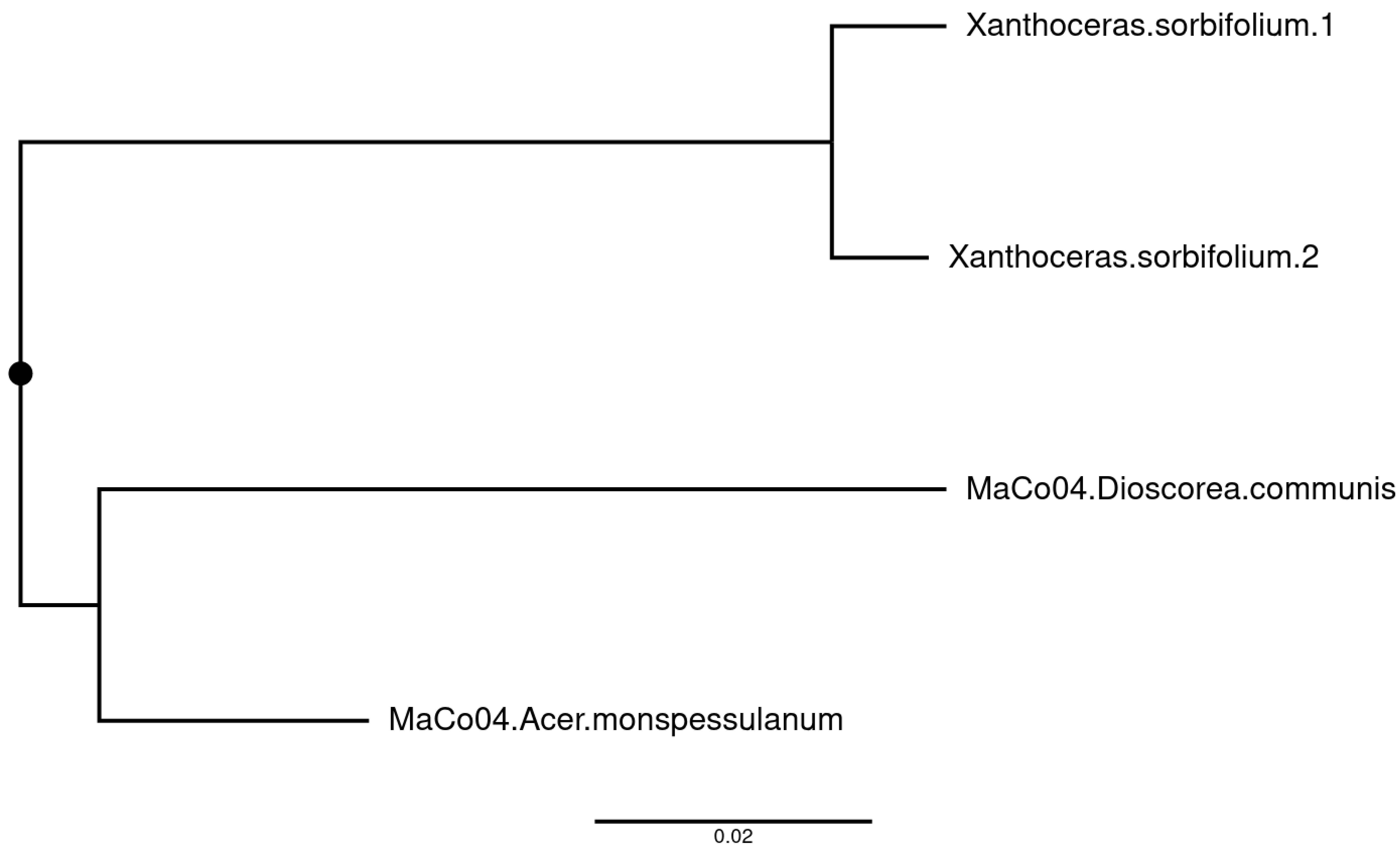

### Supplemental Figure 6

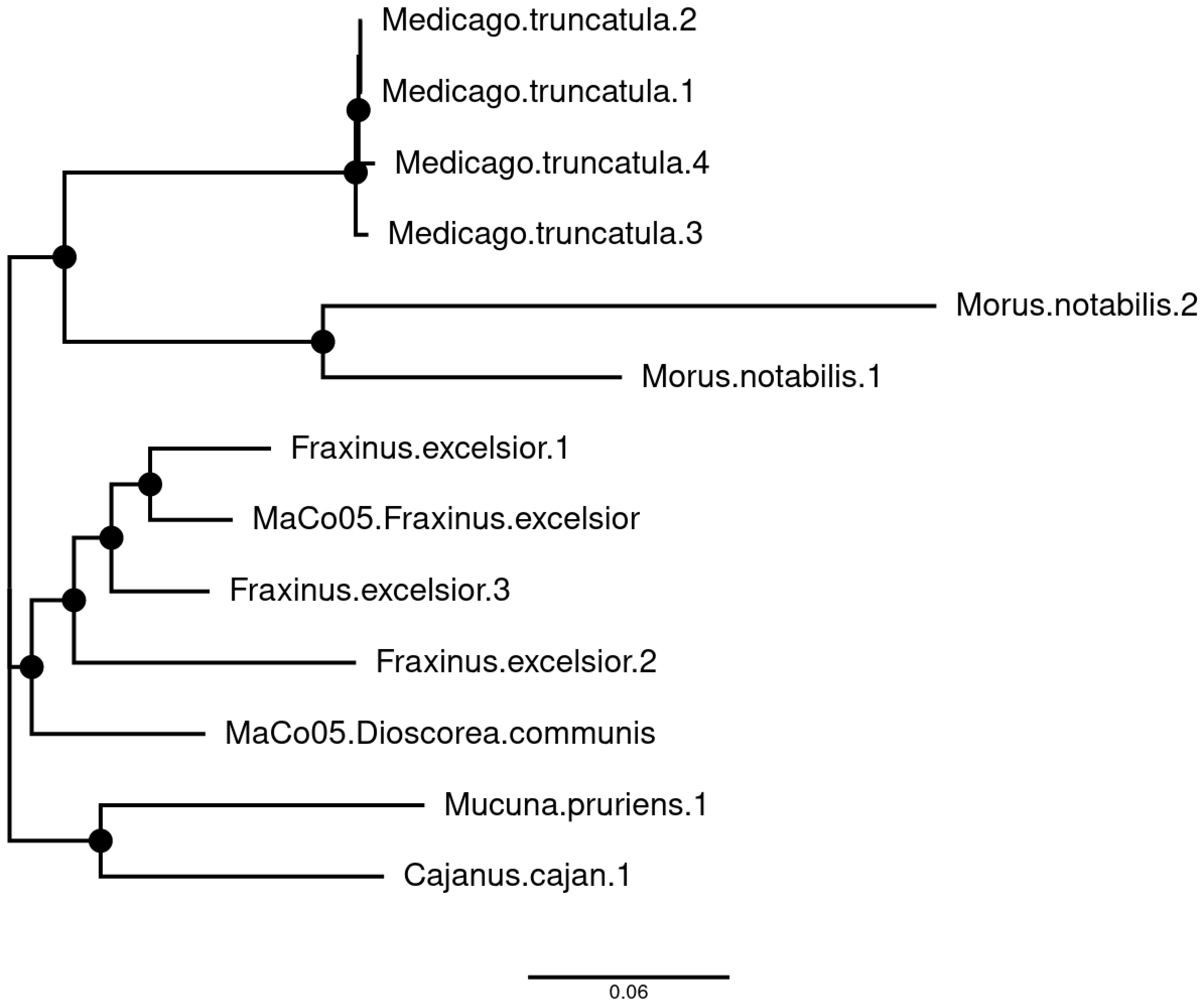

### Supplemental Figure 7

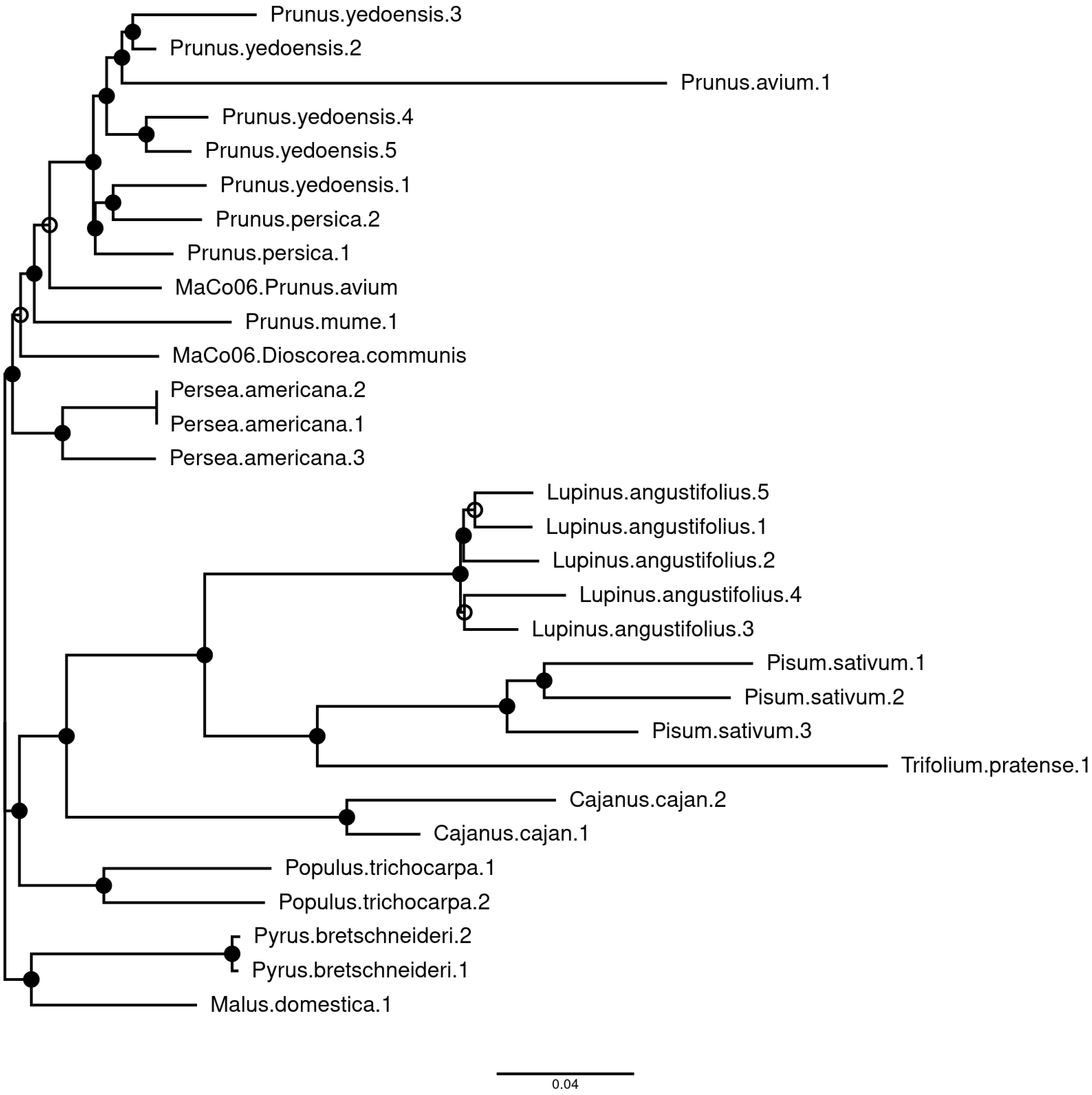

### Supplemental Figure 8

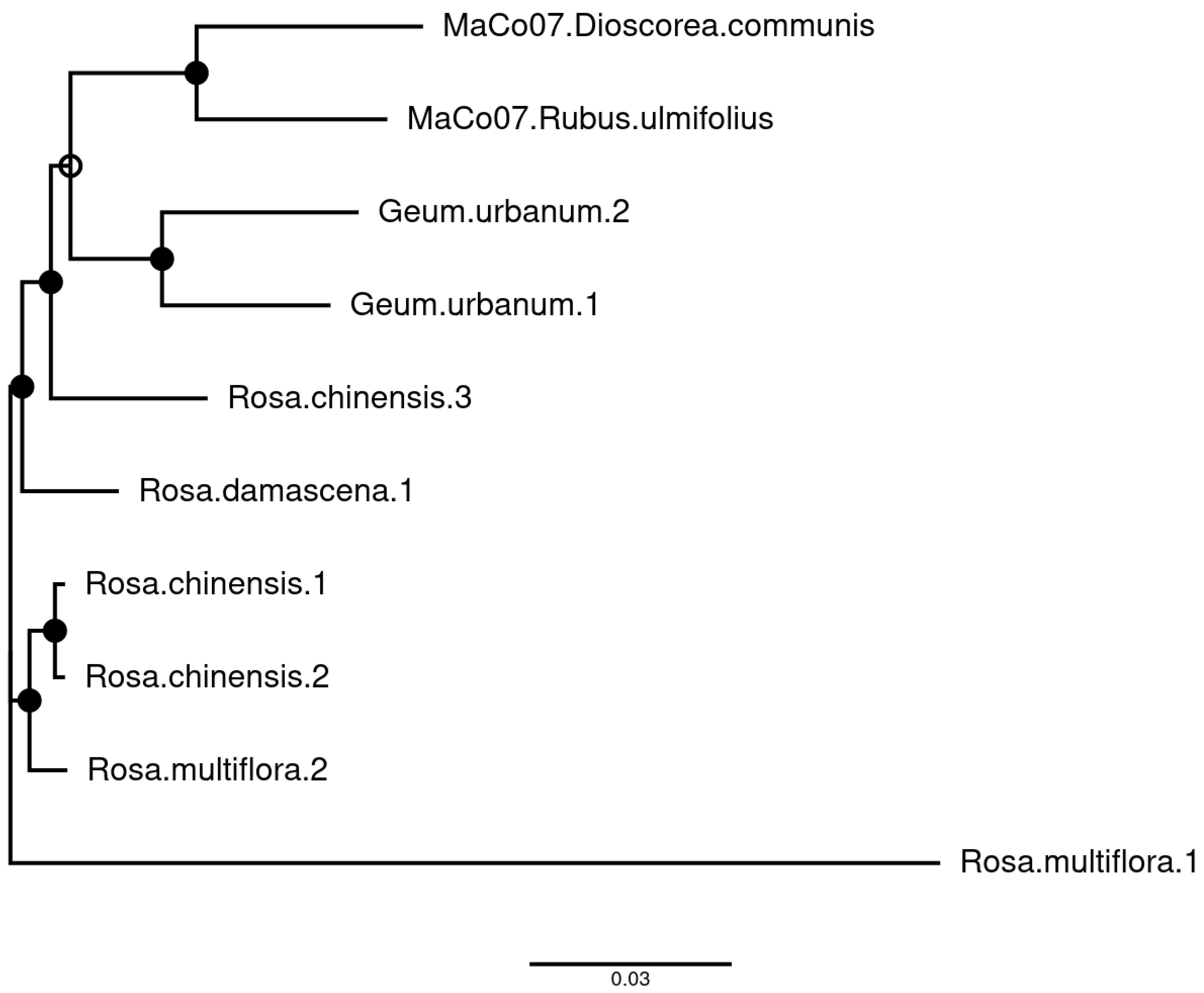

### Supplemental Figure 9

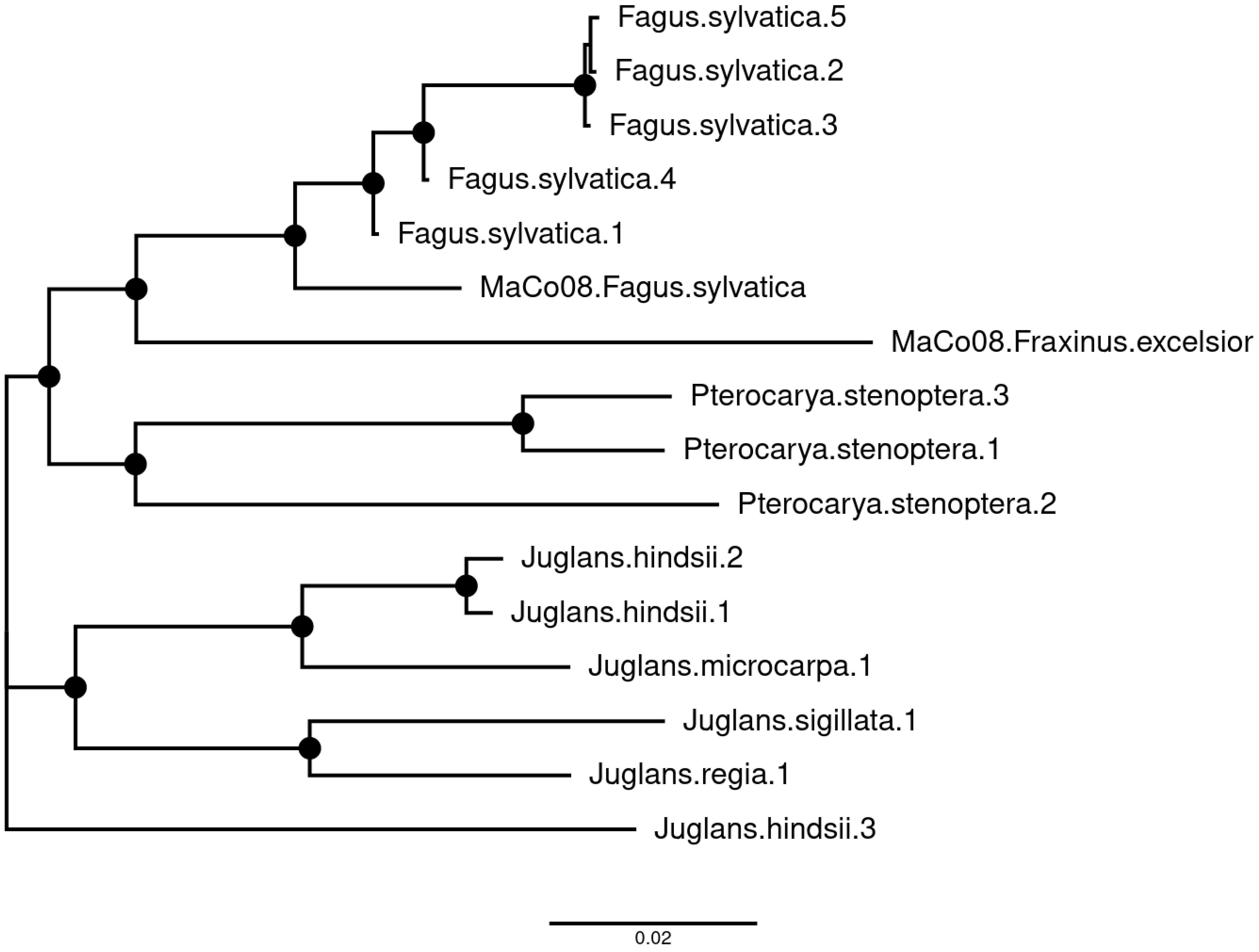

### Supplemental Figure 10

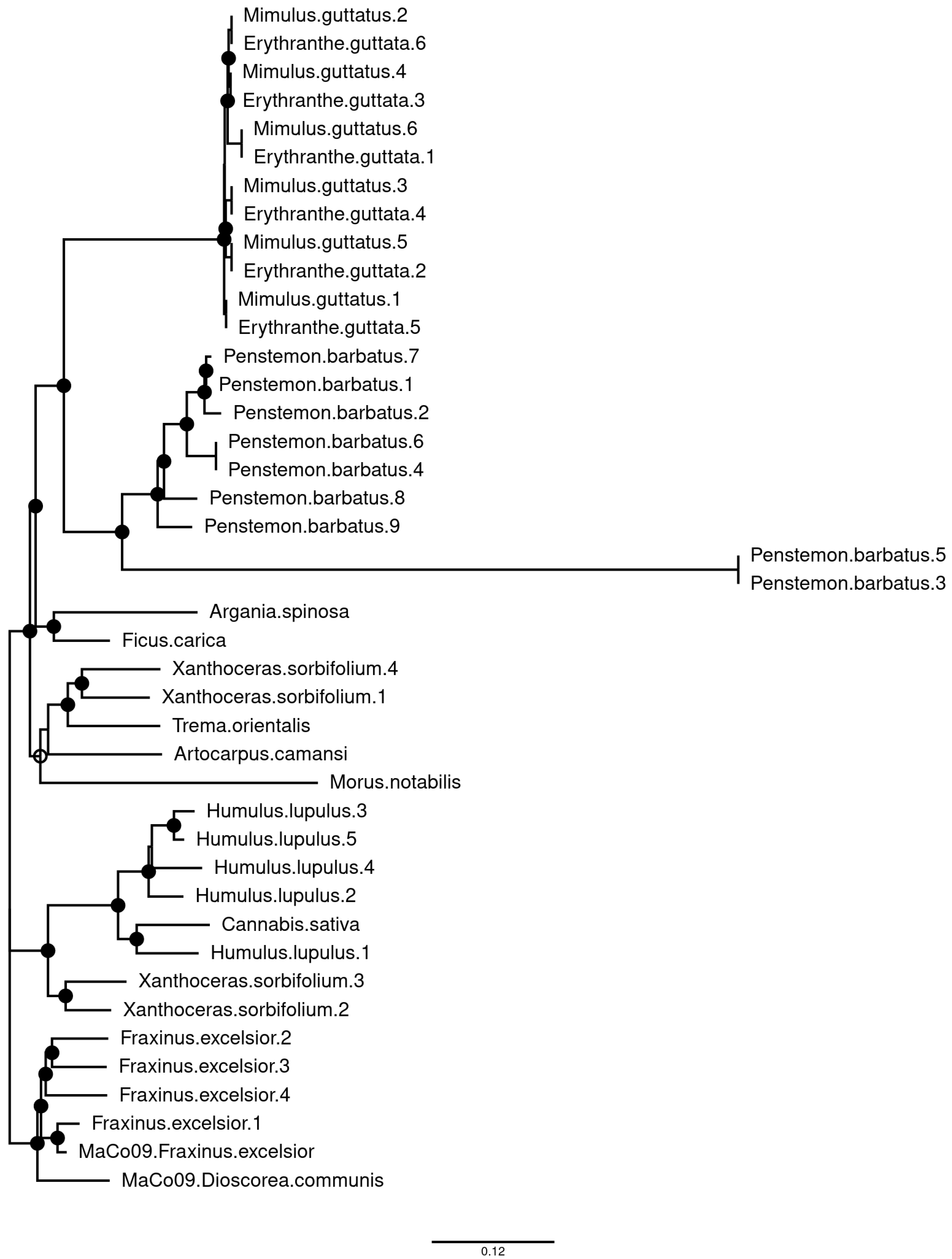

### Supplemental Figure 11

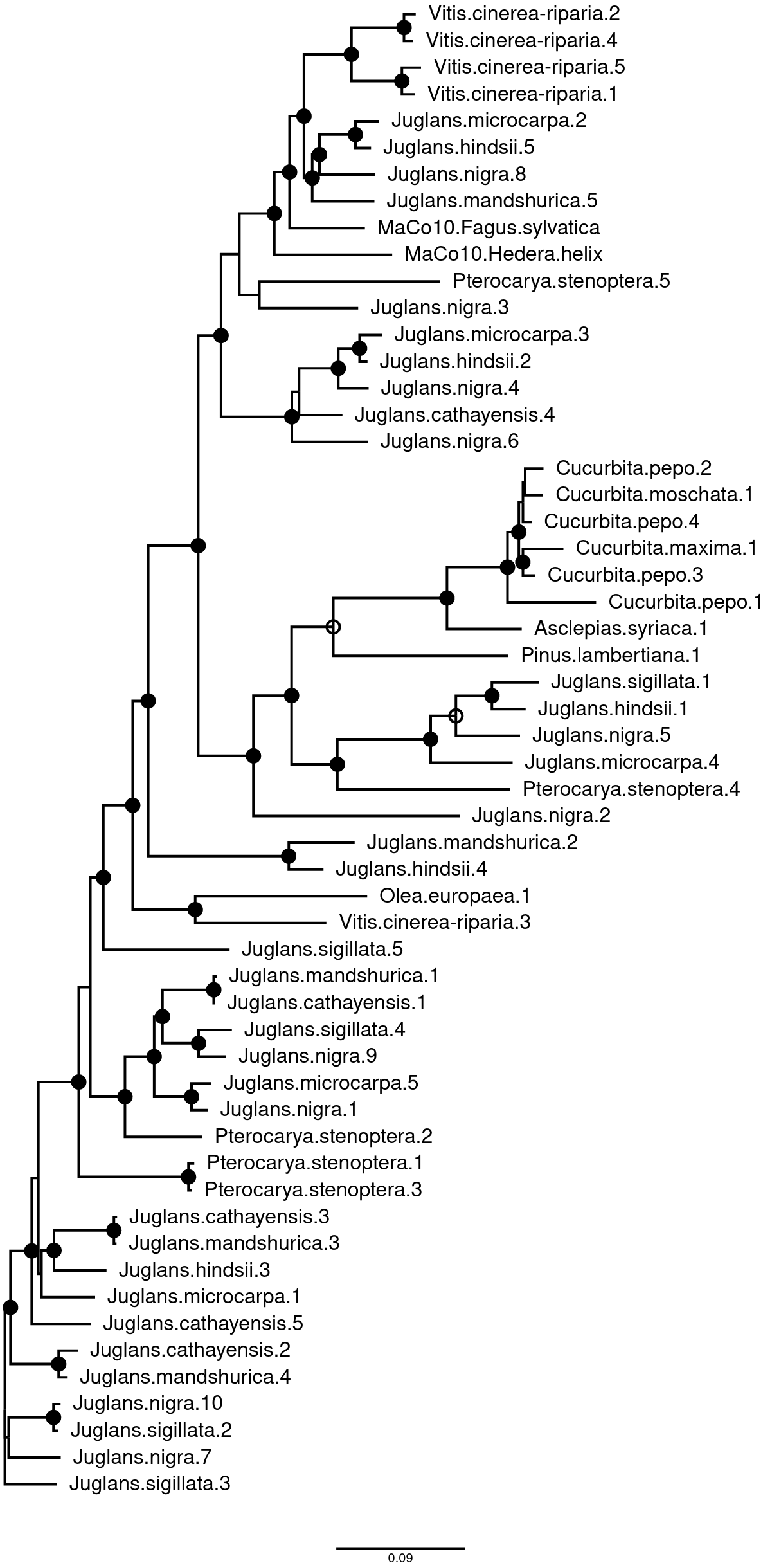

### Supplemental Figure 12

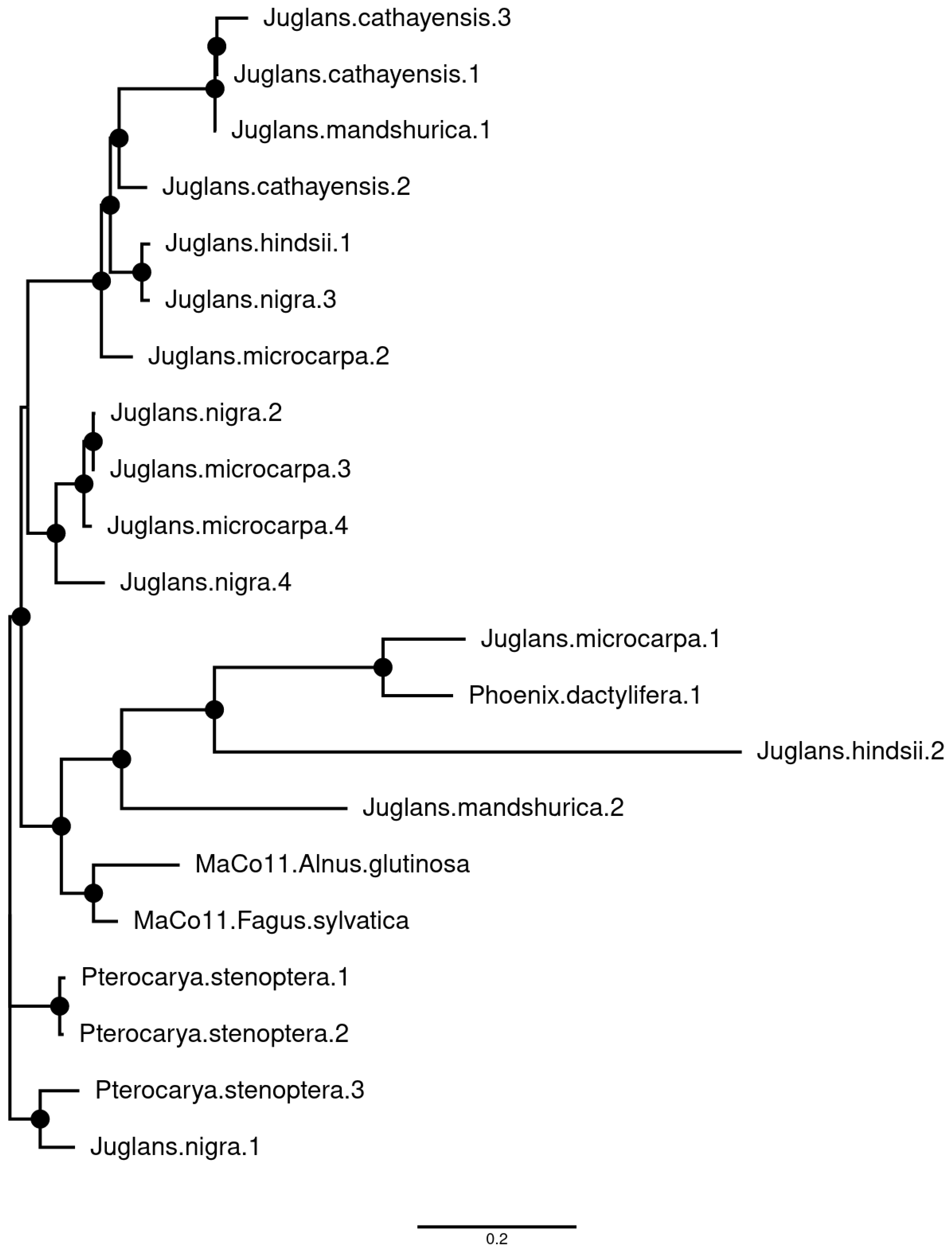

### Supplemental Figure 13

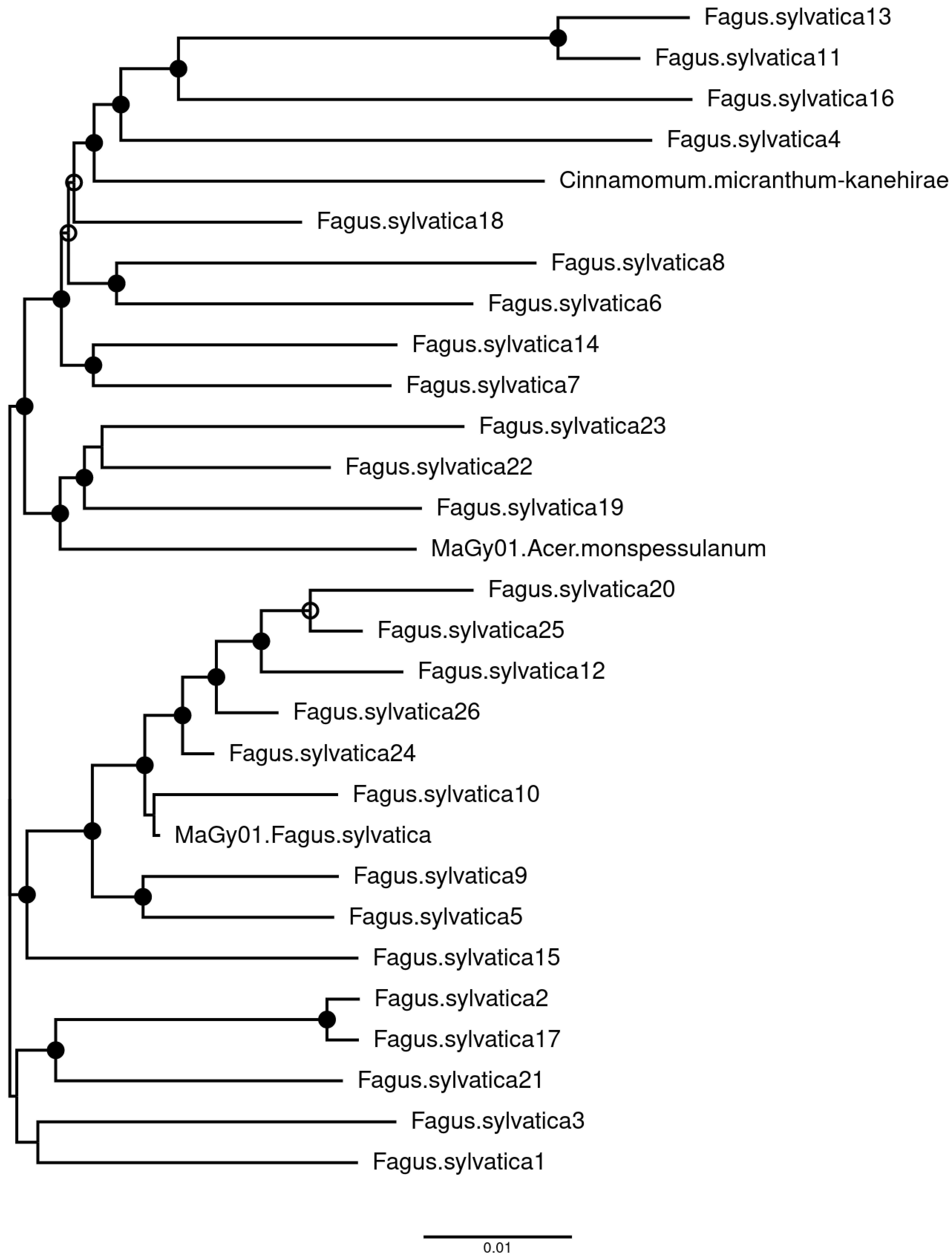

### Supplemental Figure 14

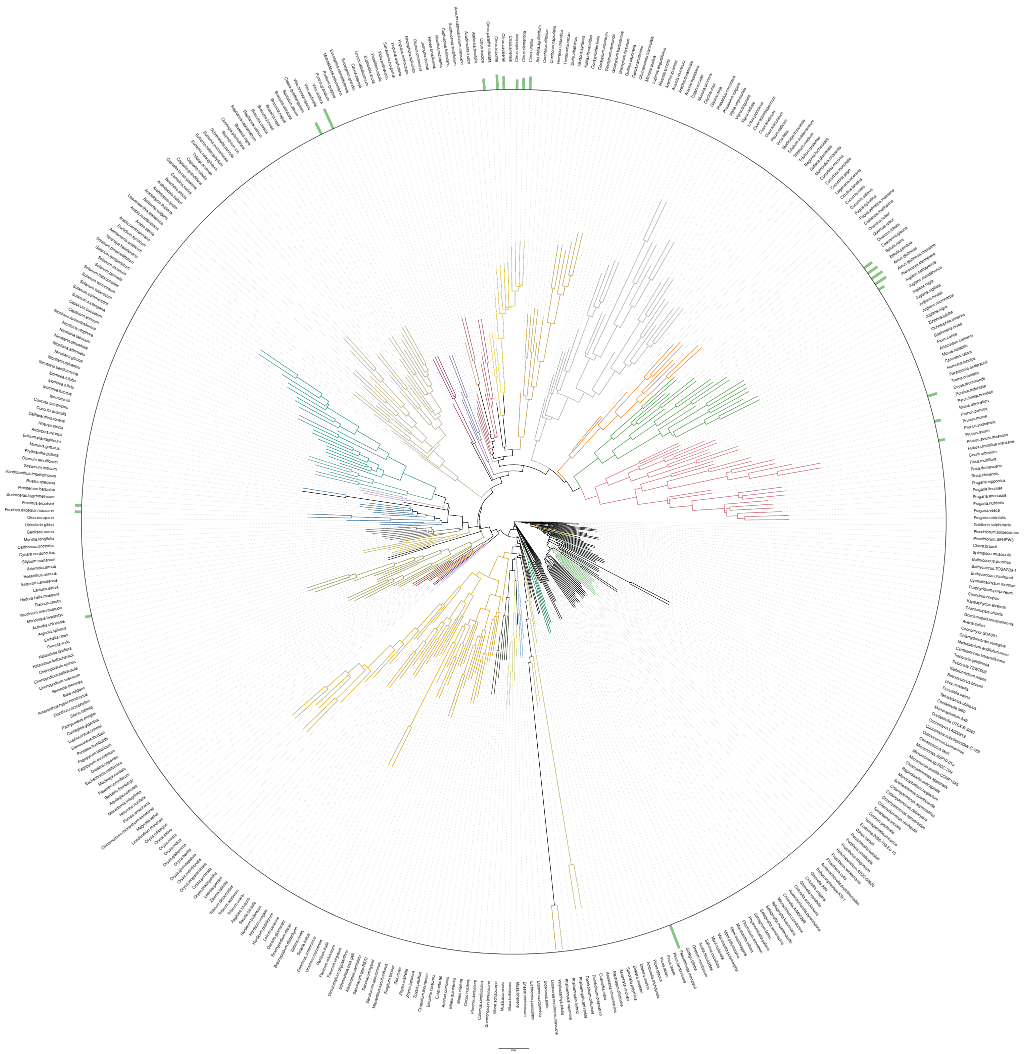

### Supplemental Figure 15

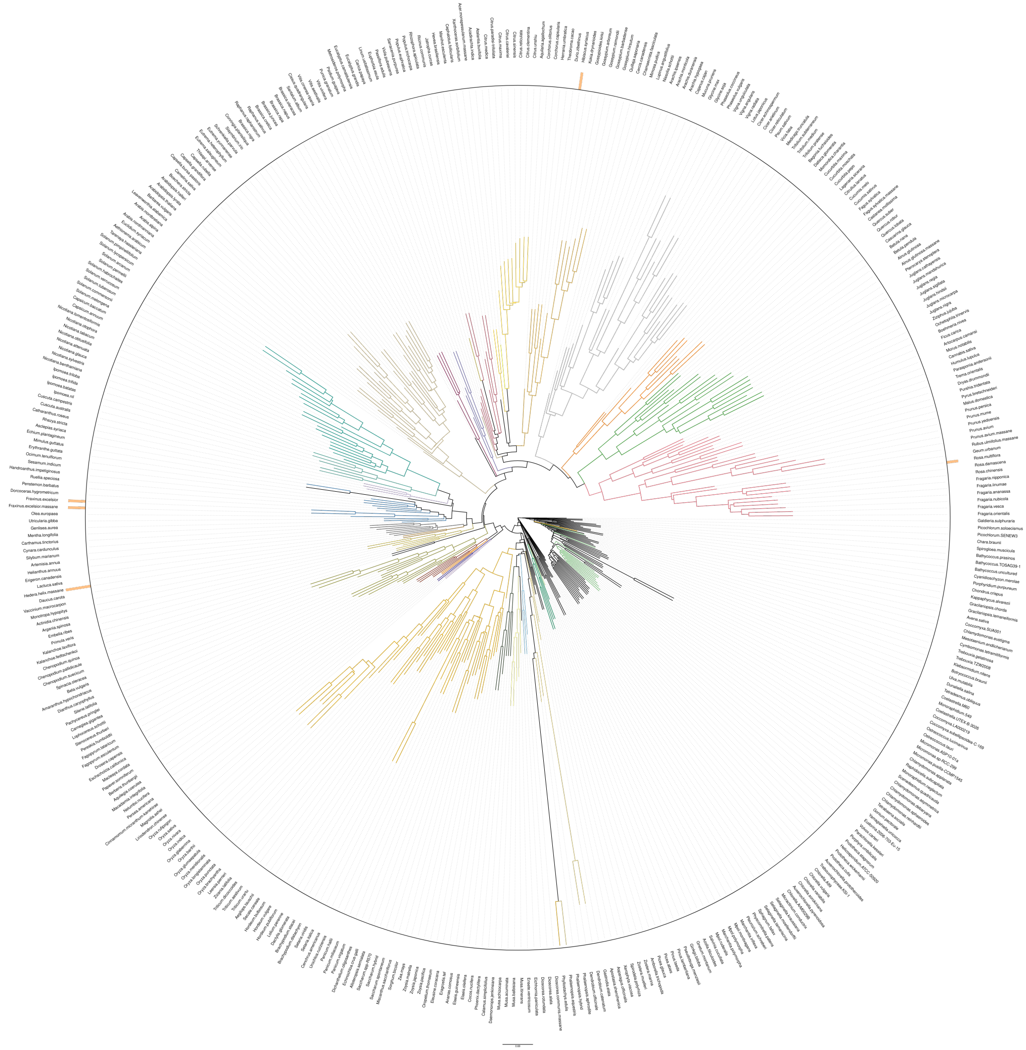

### Supplemental Figure 16

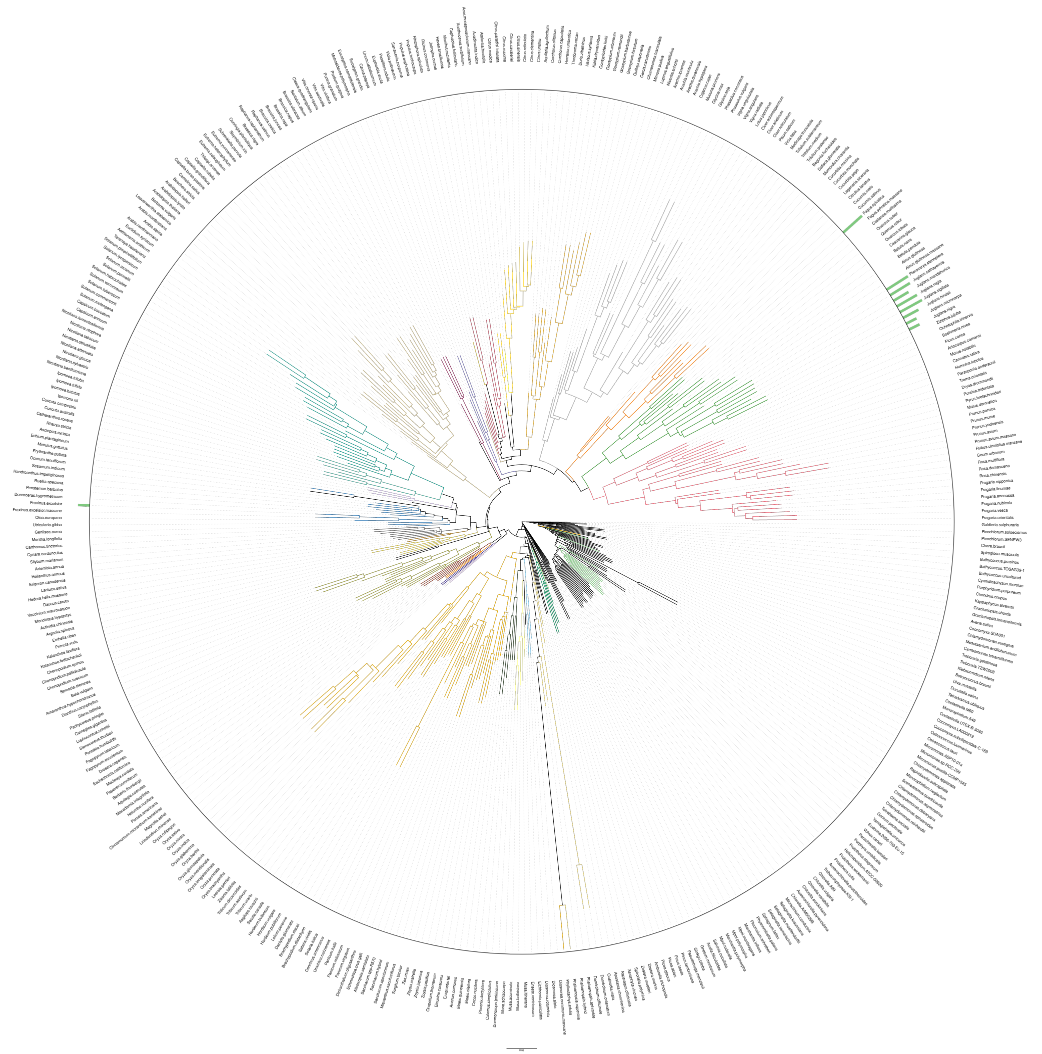

### Supplemental Figure 17

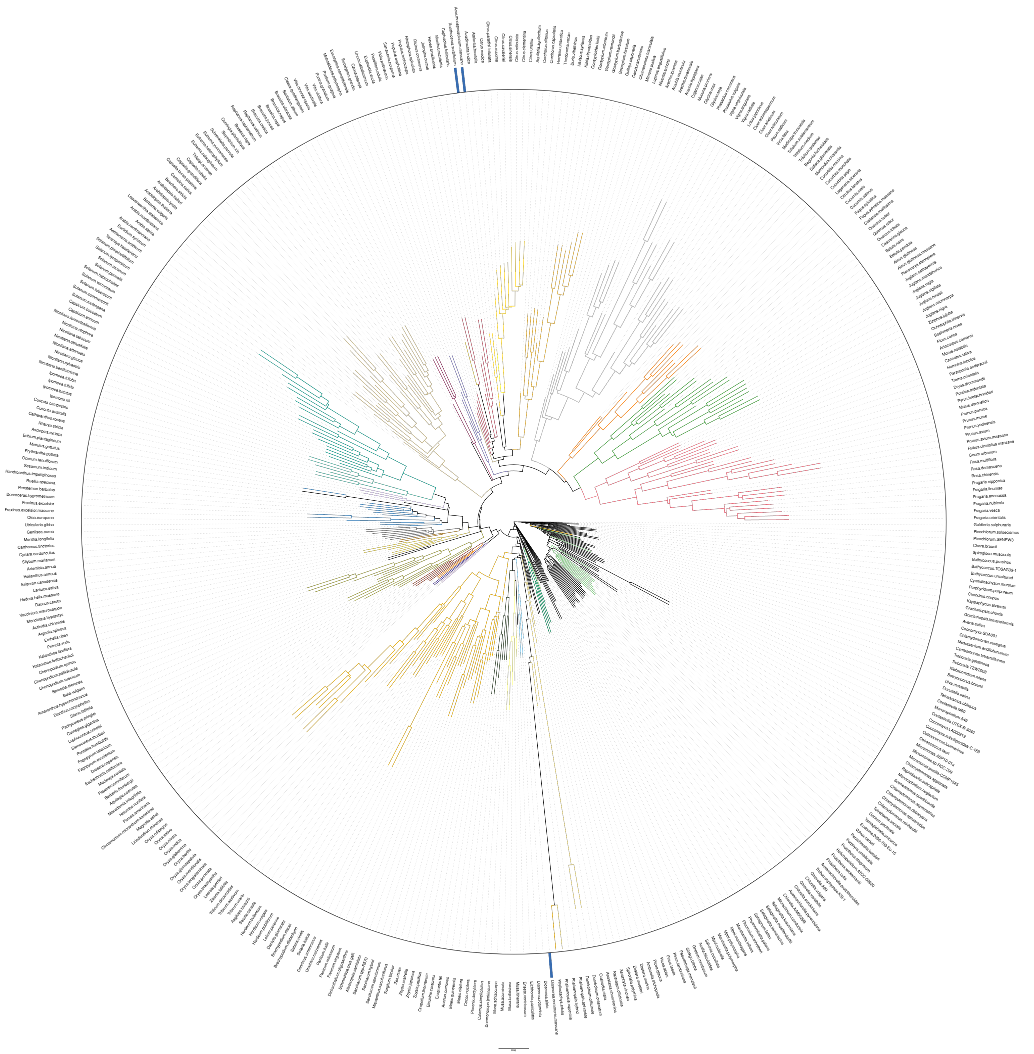

### Supplemental Figure 18

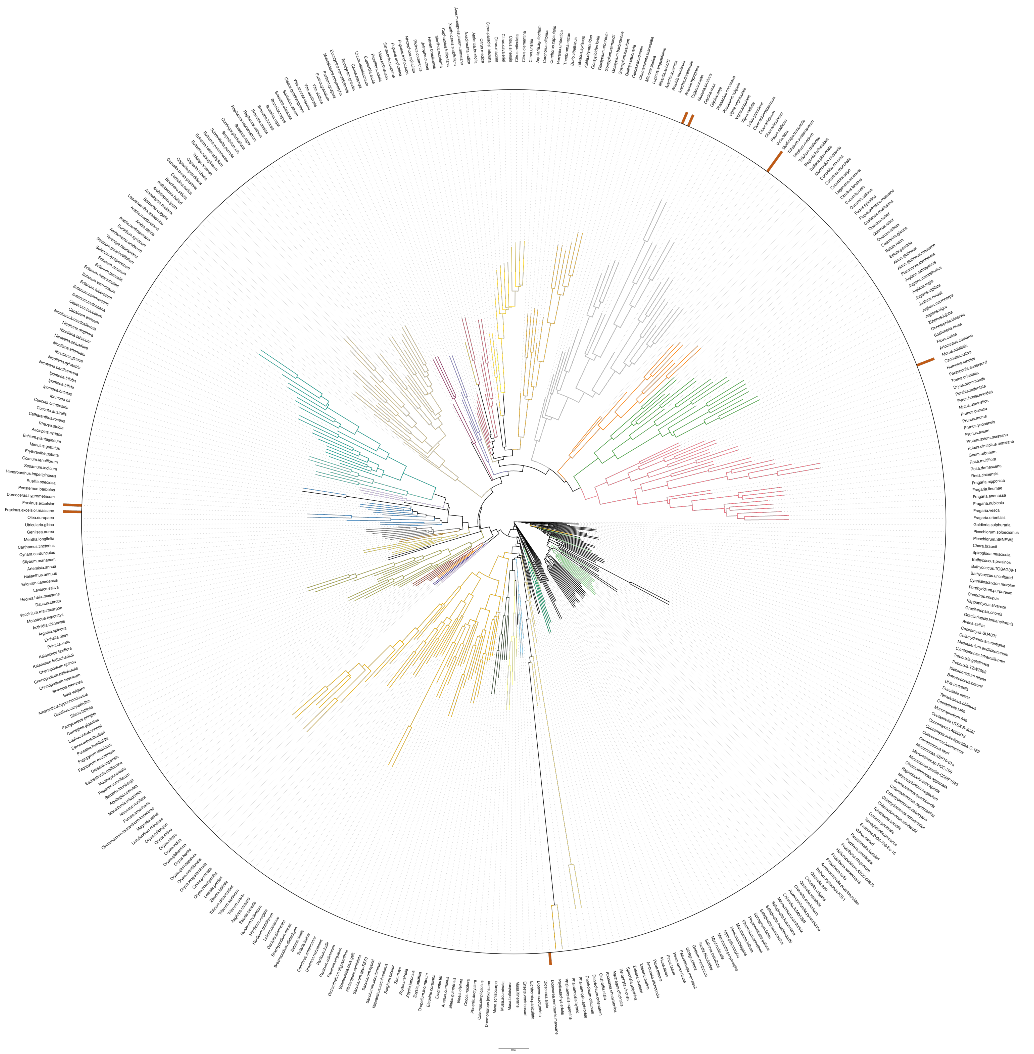

### Supplemental Figure 19

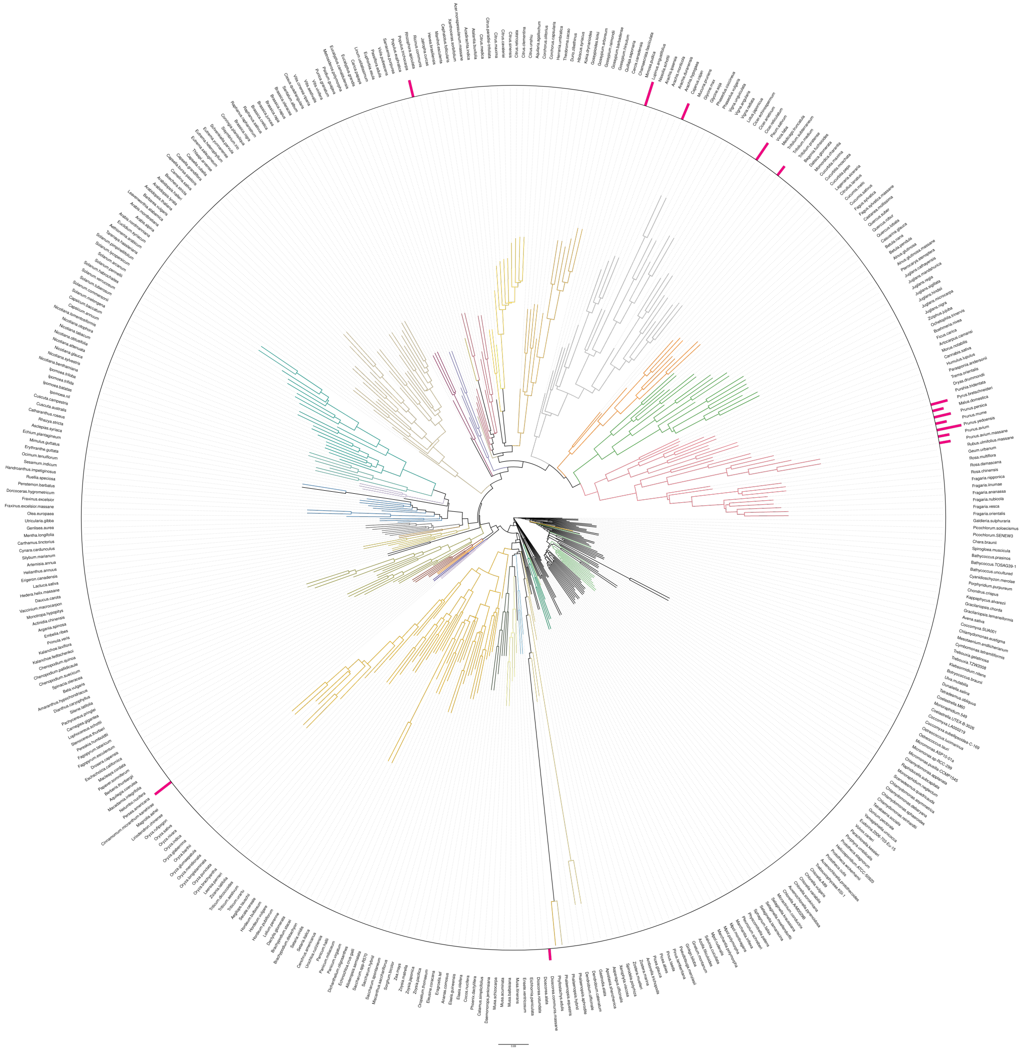

### Supplemental Figure 20

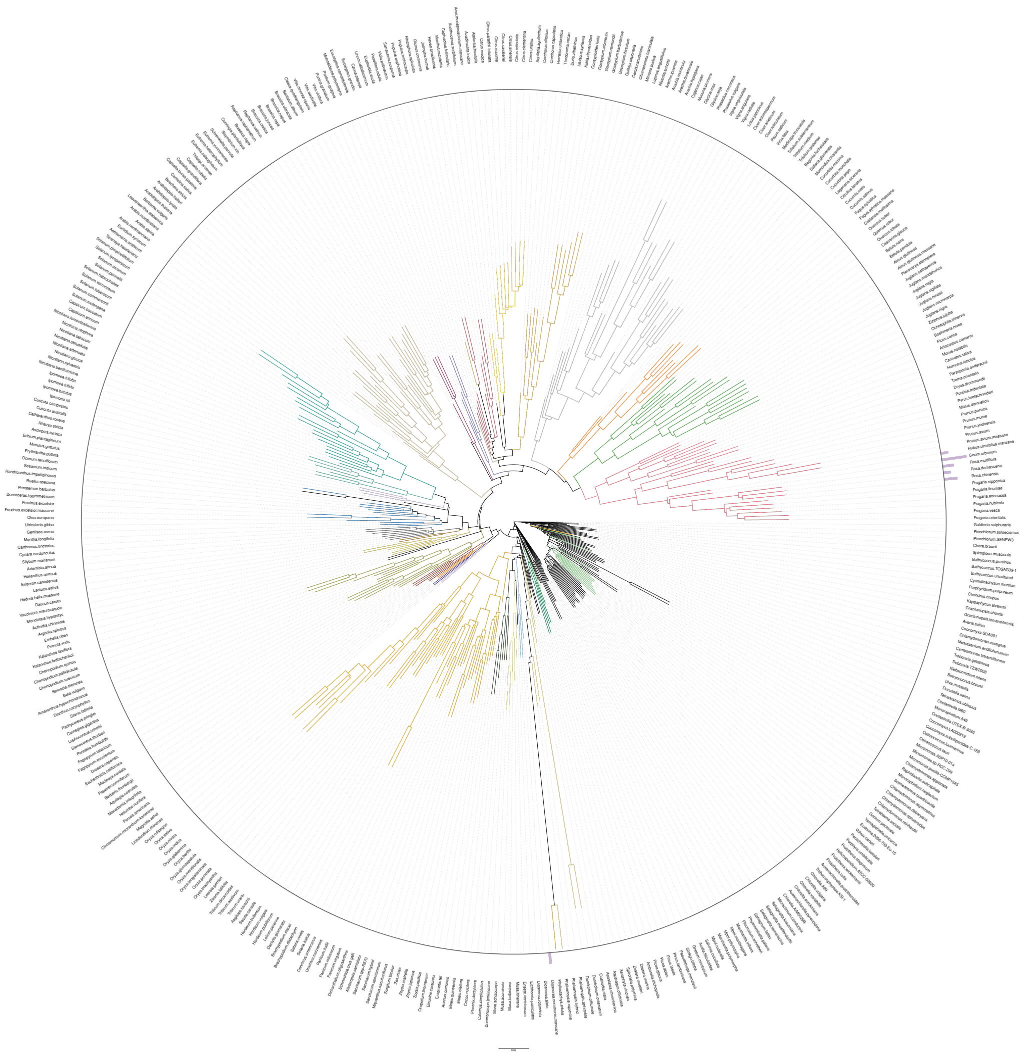

### Supplemental Figure 21

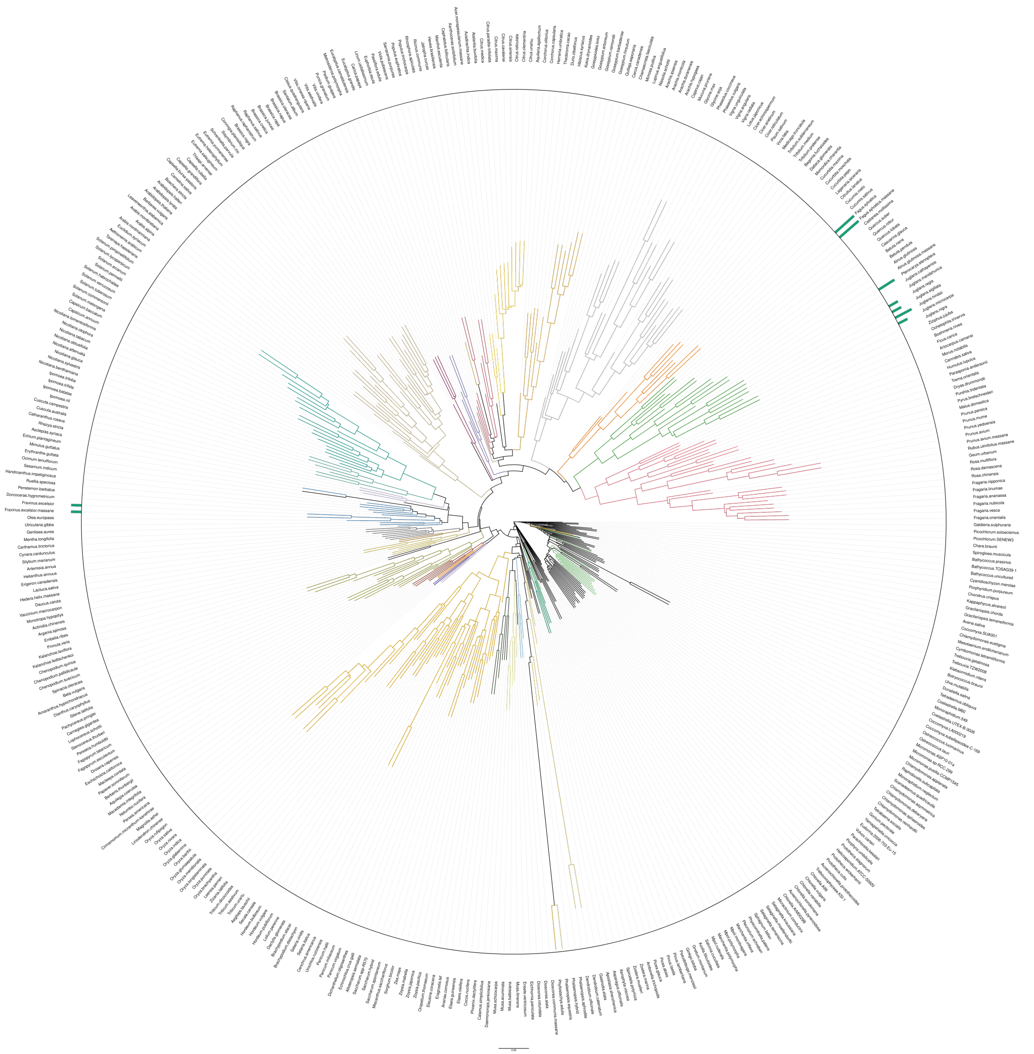

### Supplemental Figure 22

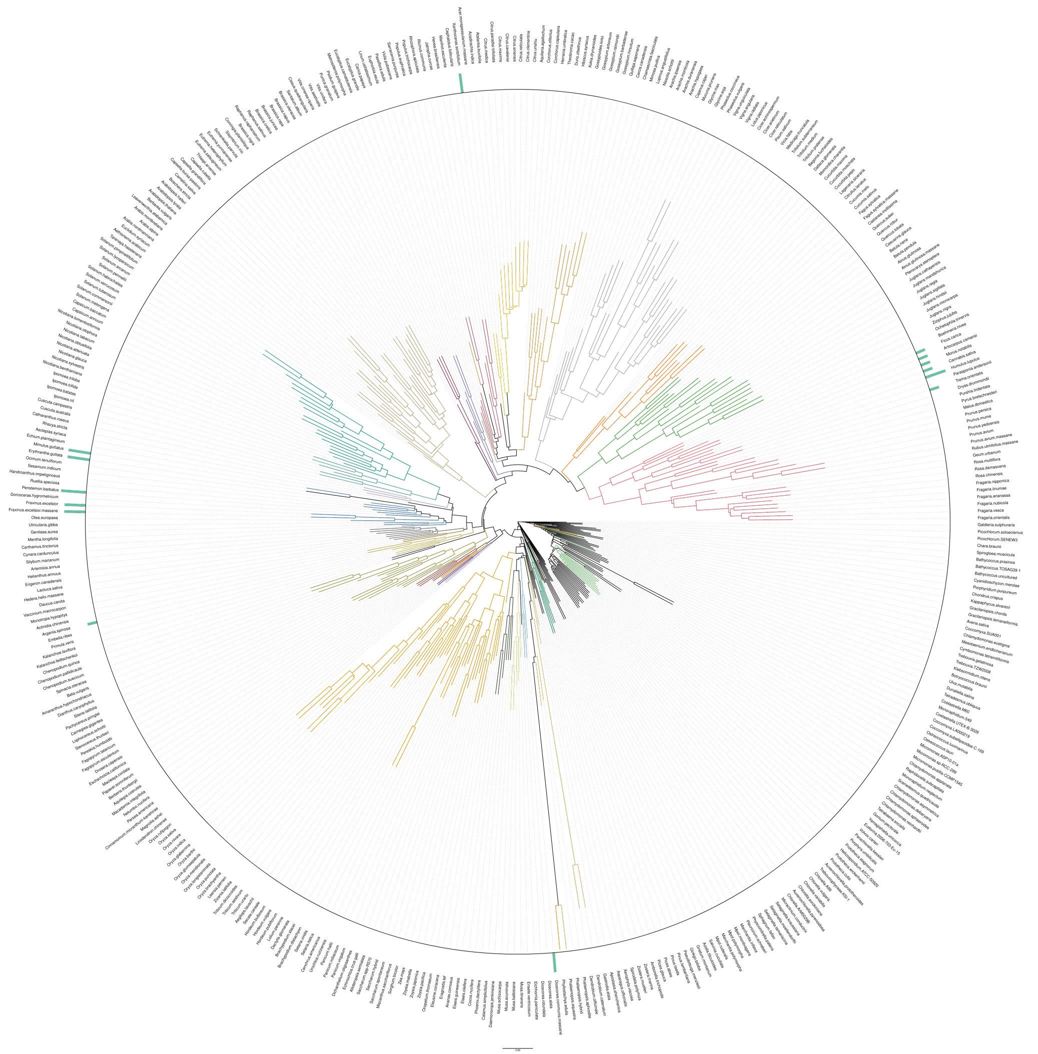

### Supplemental Figure 23

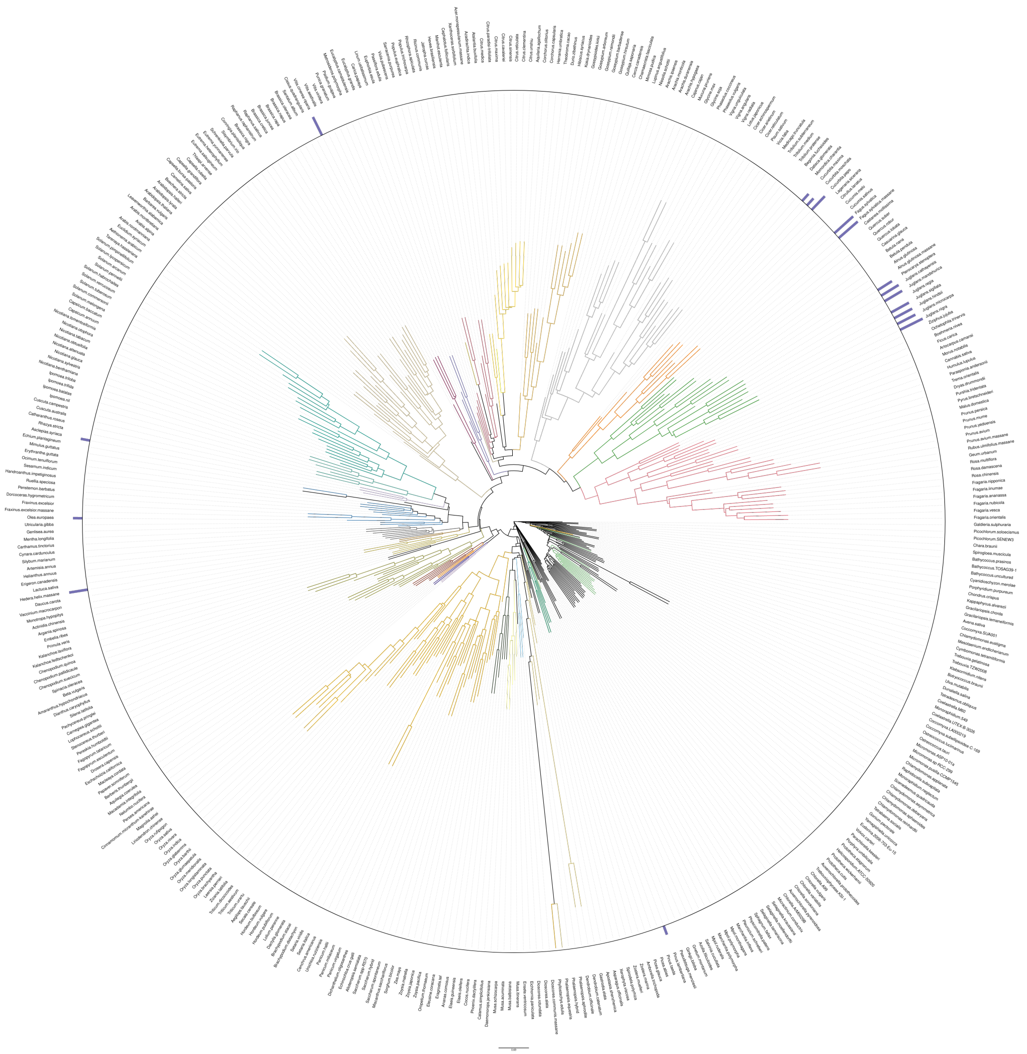

### Supplemental Figure 24

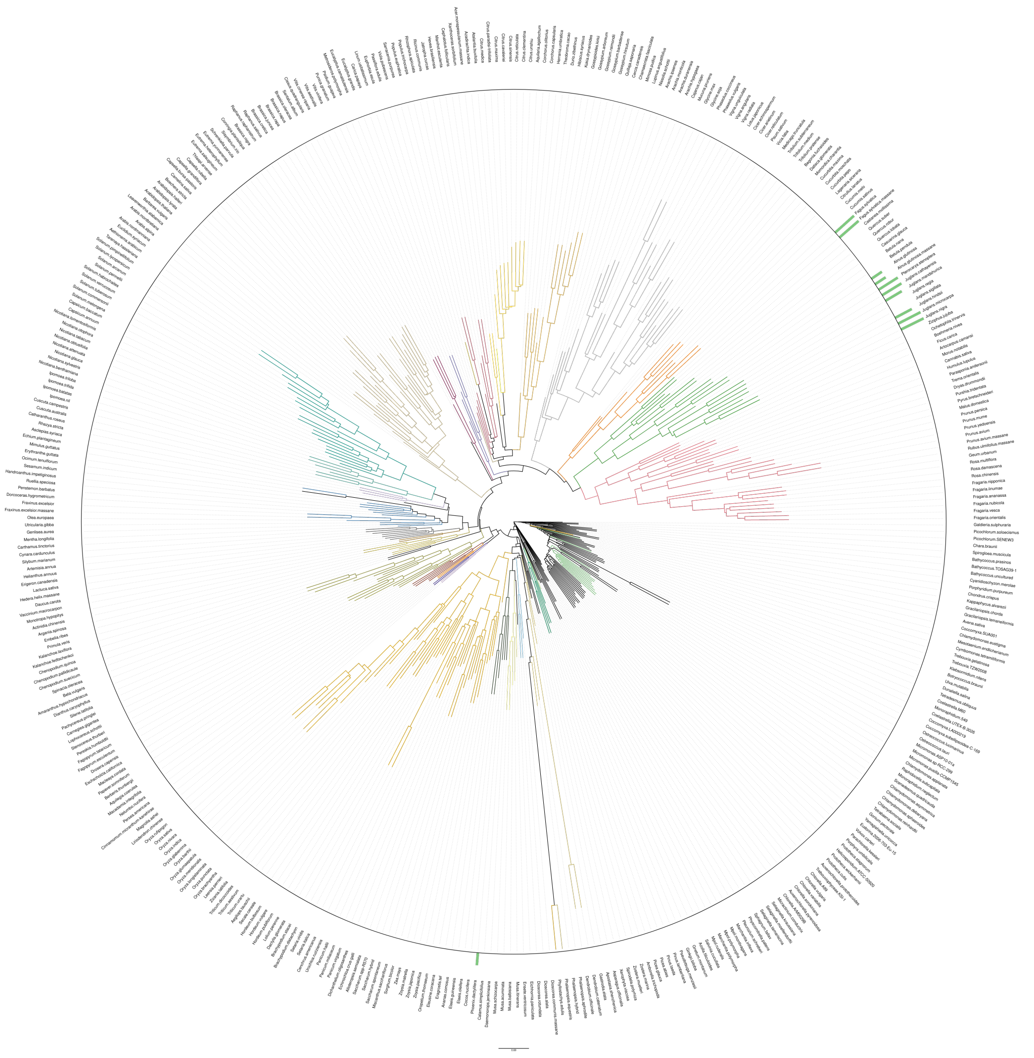

### Supplemental Figure 25

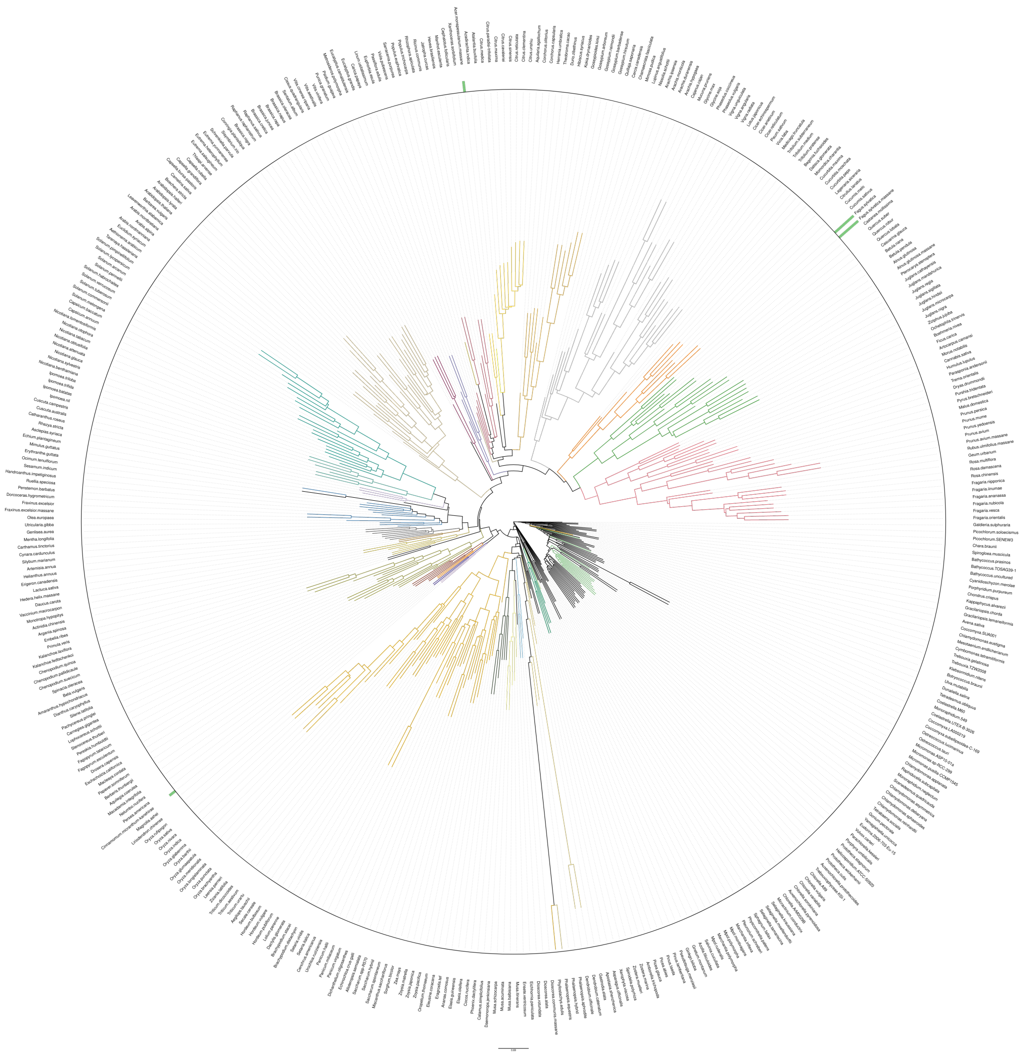

### Supplemental Figure 26

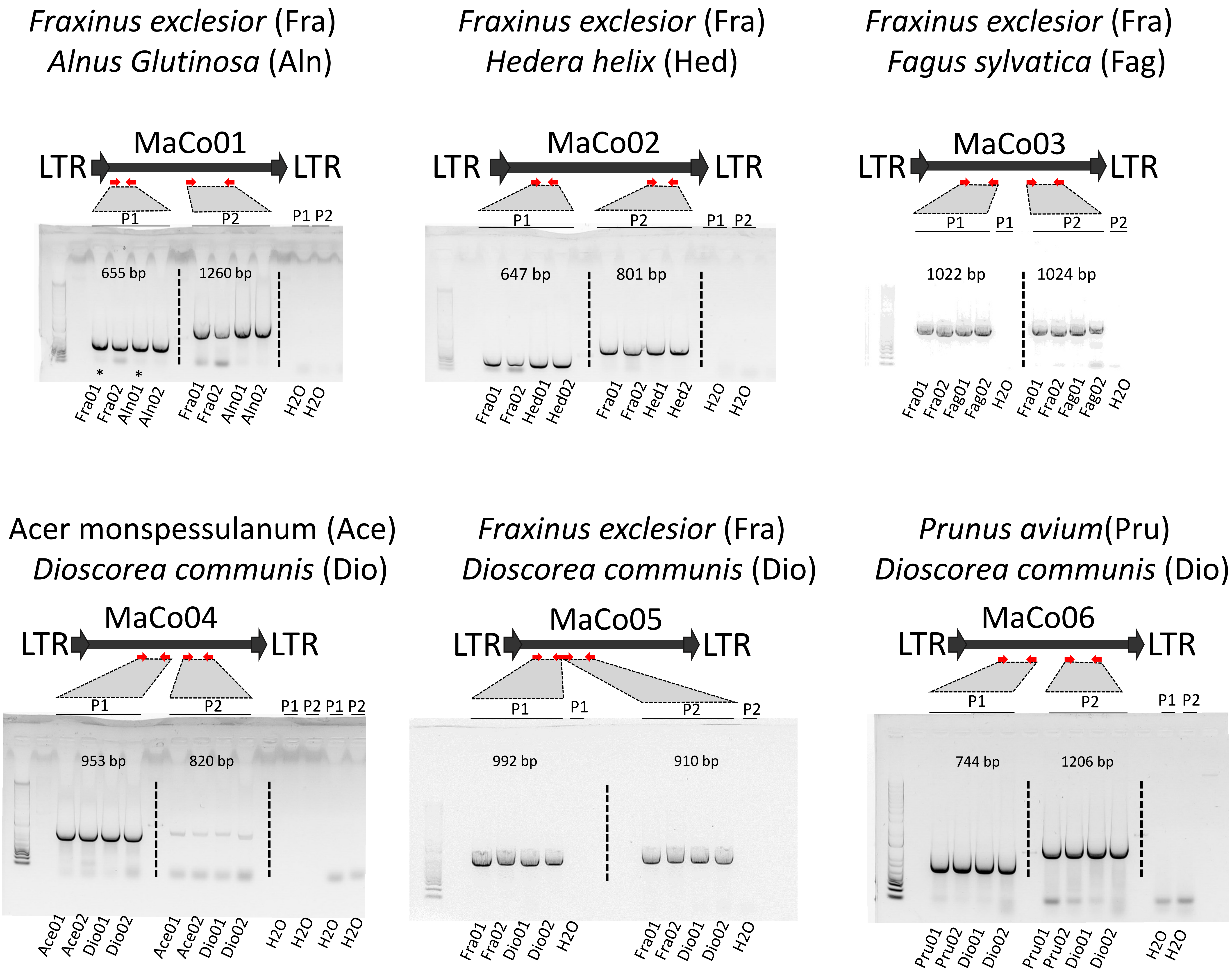

### Supplemental Figure 27

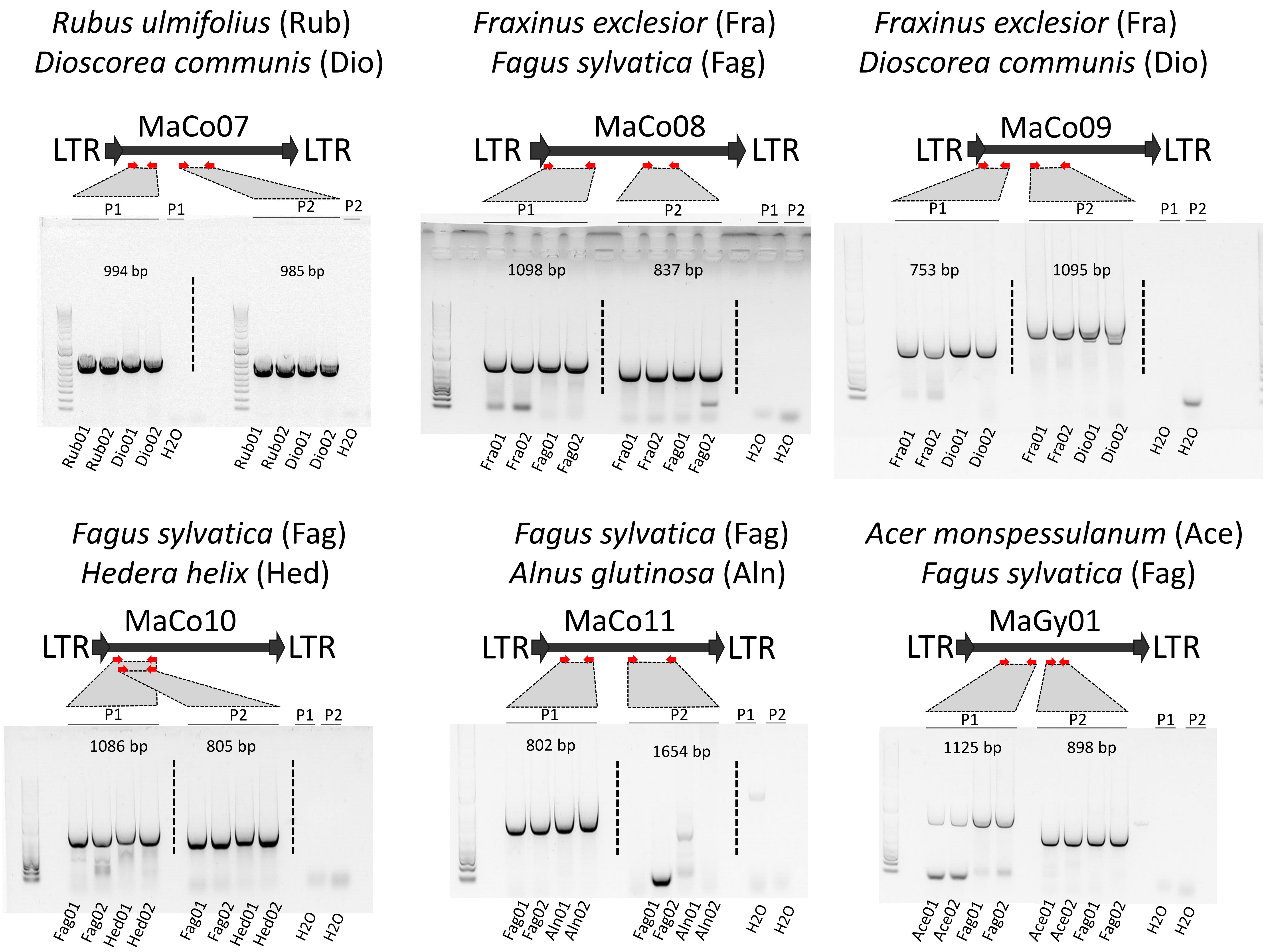

### Supplemental Figure 28

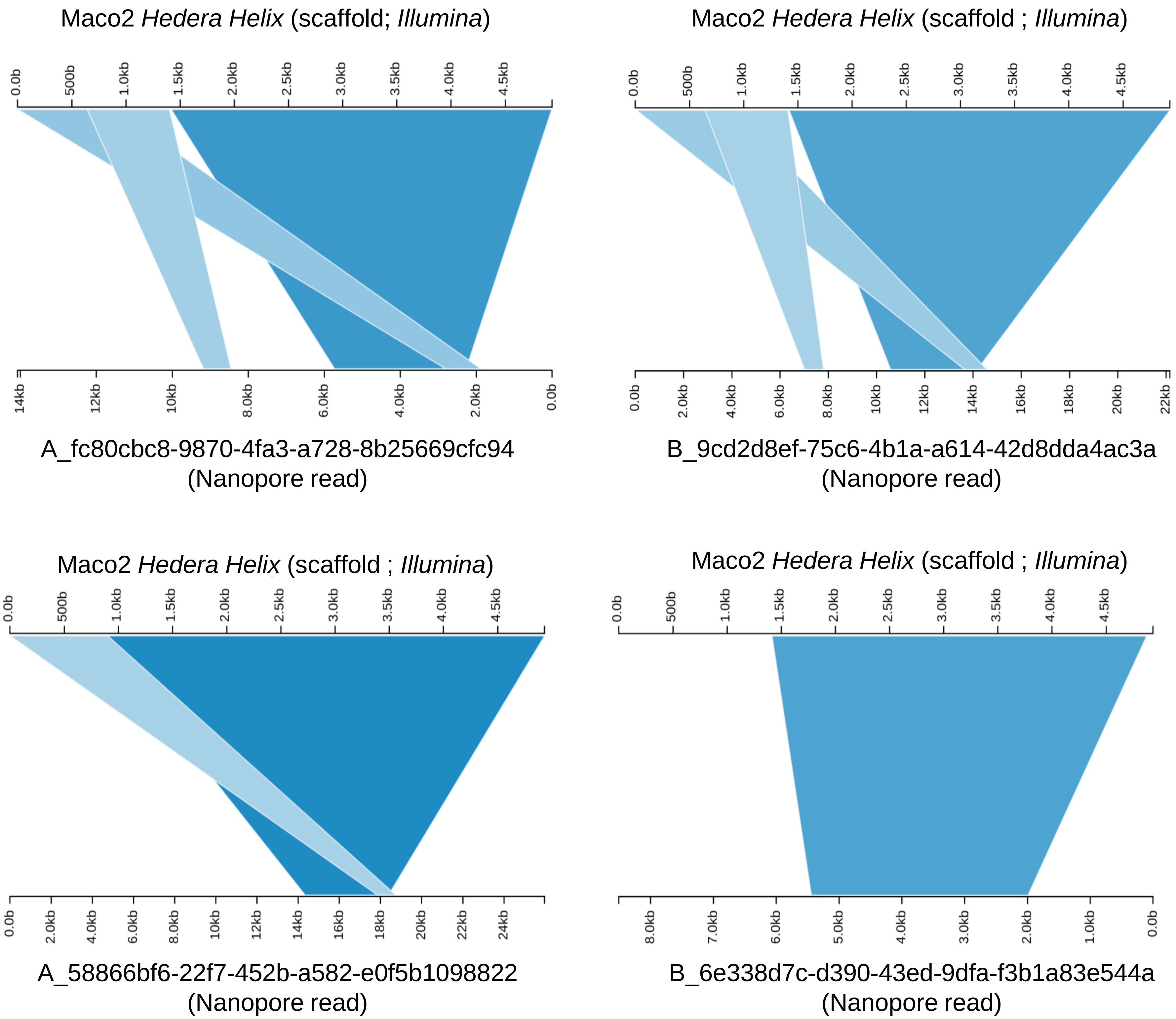

### Supplemental Figure 29

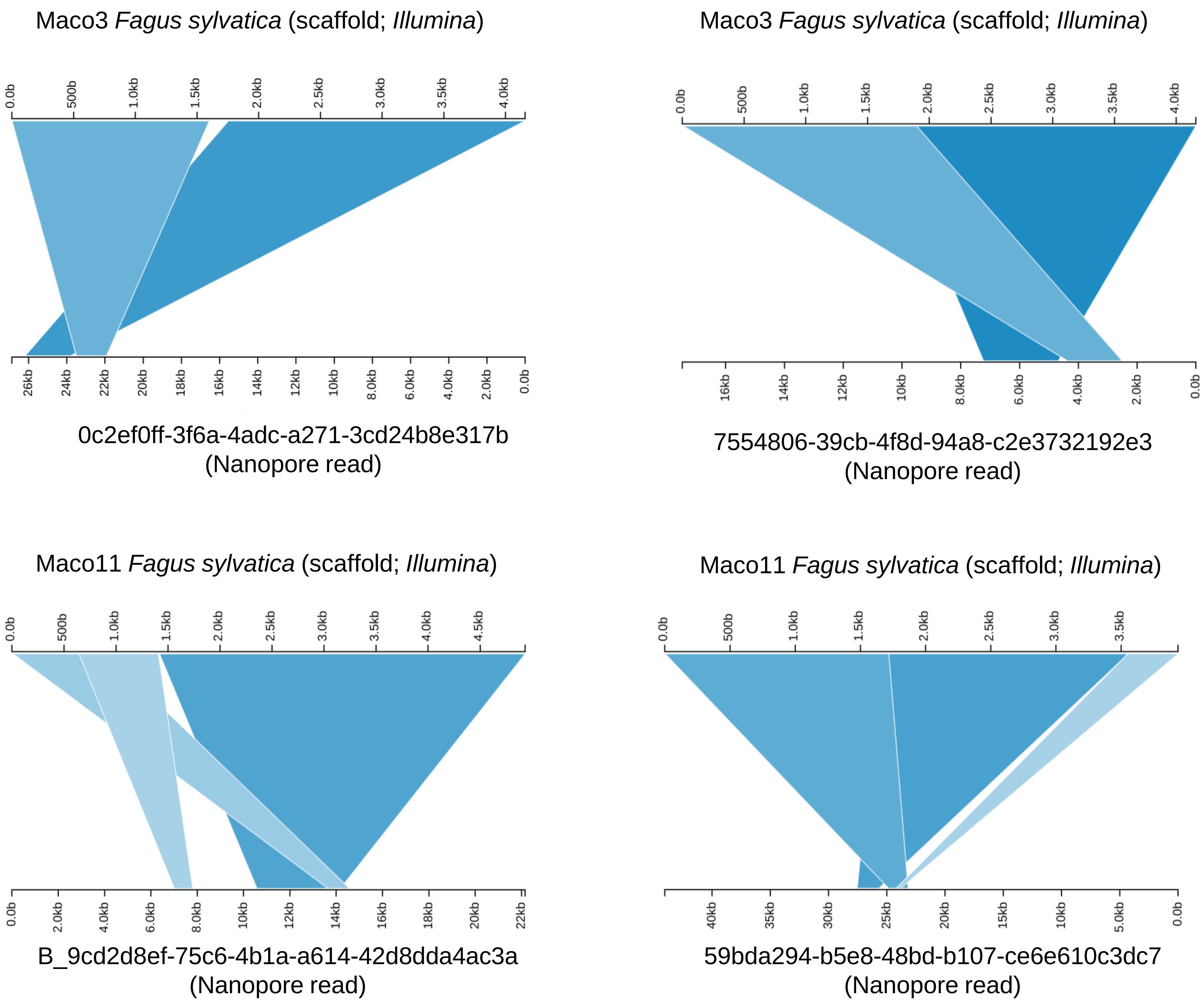
